# An expanded genetic toolkit for inducible expression and targeted gene silencing in *Rickettsia parkeri*

**DOI:** 10.1101/2024.03.15.585227

**Authors:** Jon McGinn, Annie Wen, Desmond L. Edwards, David M. Brinkley, Rebecca L. Lamason

**Affiliations:** Department of Biology, Massachusetts Institute of Technology, Cambridge, Massachusetts, USA; Department of Biological Engineering, Massachusetts Institute of Technology, Cambridge, Massachusetts, USA

## Abstract

Pathogenic species within the *Rickettsia* genus are transmitted to humans through arthropod vectors and cause a spectrum of diseases ranging from mild to life-threatening. Despite rickettsiae posing an emerging global health risk, the genetic requirements of their infectious life cycles remain poorly understood. A major hurdle toward building this understanding has been the lack of efficient tools for genetic manipulation, owing to the technical difficulties associated with their obligate intracellular nature. To this end, we implemented the Tet-On system to enable conditional gene expression in *Rickettsia parkeri*. Using Tet-On, we show inducible expression of antibiotic resistance and a fluorescent reporter. We further used this inducible promoter to screen the ability of *R. parkeri* to express four variants of the catalytically dead Cas9 (dCas9). We demonstrate that all four dCas9 variants can be expressed in *R. parkeri* and used for CRISPR interference (CRISPRi)-mediated targeted gene knockdown. We show targeted knockdown of an antibiotic resistance gene as well as the endogenous virulence factor *sca2*. Altogether, we have developed systems for inducible gene expression and CRISPRi-mediated gene knockdown for the first time in rickettsiae, laying the groundwork for more scalable, targeted mechanistic investigations into their infectious life cycles.

**IMPORTANCE:** The spotted fever group of *Rickettsia* contains vector-borne pathogenic bacteria that are neglected and emerging threats to public health. Due to the obligate intracellular nature of rickettsiae, the development of tools for genetic manipulation has been stunted, and the molecular and genetic underpinnings of their infectious lifecycle remain poorly understood. Here, we expand the genetic toolkit by introducing systems for conditional gene expression and CRISPRi-mediated gene knockdown. These systems allow for relatively easy manipulation of rickettsial gene expression. We demonstrate the effectiveness of these tools by disrupting the intracellular life cycle using CRISPRi to deplete the *sca2* virulence factor. These tools will be crucial for building a more comprehensive and detailed understanding of rickettsial biology and pathogenesis.

## INTRODUCTION

Members of the *Rickettsia* genus are obligate intracellular Gram-negative bacteria with a broad host range^1–3^. Several *Rickettsia* species are neglected human pathogens transmitted by arthropod vectors like ticks, mites, fleas, and lice, and some are among the oldest known vector-borne pathogens^4^. Still, we know little about the genetic and molecular requirements of their infectious lifecycle. This knowledge gap is largely due to the challenges associated with studying obligate intracellular bacteria, including the lack of a modern toolkit to perform targeted genetic manipulation in these pathogens^3,5,6^.

Over the course of their evolution as obligate intracellular bacteria, rickettsiae have undergone drastic genome reduction and rearrangement, giving rise to small, streamlined genomes^1,7^. Rickettsial genomes typically contain fewer than 1500 predicted coding sequences in their 1.1-1.5 Mb genomes^7^. Despite these small genomes, only a small fraction of rickettsial genes have been studied in detail, and even fewer have been directly studied in mutant strains of *Rickettsia*^1,5,6,8,9^. As is the case in other obligate intracellular bacteria, the development of tools for genetic manipulation in rickettsiae has lagged behind many other model bacteria^5^. Plasmid-driven transposon mutagenesis was only first reported in 2007^10^ and later adopted by others in the field^8,9,11,12^. It was not until 2011 that a shuttle vector was generated for use in rickettsiae^13^, leading to the first genetic complementation of a mutant in 2016^14^. Reports of targeted genetic knockouts or silencing in rickettsiae have also emerged, using approaches based on allelic exchange^15,16^, group II intron mutagenesis^17^, and peptide nucleic acids^18^. However, these approaches are not always amenable to studying essential genes and are often low throughput. Additionally, no systems for conditional gene expression have been reported in rickettsiae. Thus, easy and scalable methods for targeted control of rickettsial gene expression would greatly advance the field’s ability to carry out detailed mechanistic analyses of the rickettsial infectious life cycle.

In the last decade, CRISPR-based tools have enabled genetic manipulation in many previously intractable organisms^19^. One such tool is CRISPR interference (CRISPRi), which relies on a catalytically dead mutant of Cas9 (dCas9) to reversibly knock down genes of interest by physically blocking transcription initiation and/or elongation^20,21^. dCas9 is directed to genomic loci of interest via sequence homology with a guide RNA (gRNA). This homology search between gRNA and the genome is licensed by direct interactions between the dCas9 protein and a protospacer adjacent motif (PAM)^22^. Different variants of dCas9 recognize different PAMs^23^, meaning that each dCas9 has a different repertoire of possible guide sequences that can be used to target a genome of interest. CRISPRi has been used for efficient and scalable gene knockdown in a wide variety of bacteria including *Mycobacterium tuberculosis*^24^, *Caulobacter crescentus*^25^, *Chlamydia trachomatis*^26^, and *Coxiella burnetii*^27,28^.

Here, we expand the rickettsial genetic toolkit by introducing systems for conditional gene expression and targeted gene knockdown via CRISPRi in *Rickettsia parkeri*. We demonstrate the feasibility of conditional gene expression through inducible expression of an antibiotic-resistance gene and a fluorescent reporter gene using the Tet-ON system. We were subsequently able to use this conditional expression system to express four variants of dCas9 in *R. parkeri*, all of which enabled knockdown of an antibiotic-resistance gene. We further show CRISPRi-mediated knockdown of the endogenous virulence gene *sca2*. Altogether, this work greatly expands the arsenal of genetic tools for rickettsiae, opening new avenues for mechanistic investigations into rickettsial biology and pathogenesis.

## RESULTS

### Development of an inducible promoter system for Rickettsia

We first set out to build a system for conditional gene expression in *Rickettsia parkeri*. We chose the Tet-On system developed from the Tn*10* transposon of *Escherichia coli*^29,30^, based on the membrane-permeability of tetracycline derivatives in mammalian host cells^31^ and previous success of implementing a tetracycline-inducible promoter in *Chlamydia trachomatis*^32^. To assess feasibility, we first needed to determine the viability of *R. parkeri* upon exposure to anhydrotetracycline (aTc), which is widely used as the inducer of Tet-On^33^. To measure aTc toxicity, we infected Vero host cells with *R. parkeri* and added various concentrations of aTc at the time of infection. We then imaged and quantified plaque formation at five days post-infection (dpi). Plaques were observed at aTc concentrations from 0.1 to 250 ng/mL, indicating successful *R. parkeri* infection at these concentrations (Fig. 1A). At 100 and 250 ng/mL, plaque numbers began to trend downward but did not reach statistical significance, suggesting slight aTc toxicity may occur at these concentrations. At 500 ng/mL, no plaques were formed, indicating sensitivity of *R. parkeri* to higher concentrations of aTc, similar to what was observed with *C. trachomatis*^32^. These results demonstrate that aTc is well tolerated during infection, indicating that the Tet-On system may be suitable for inducible expression in *R. parkeri*.

**Figure 1.**
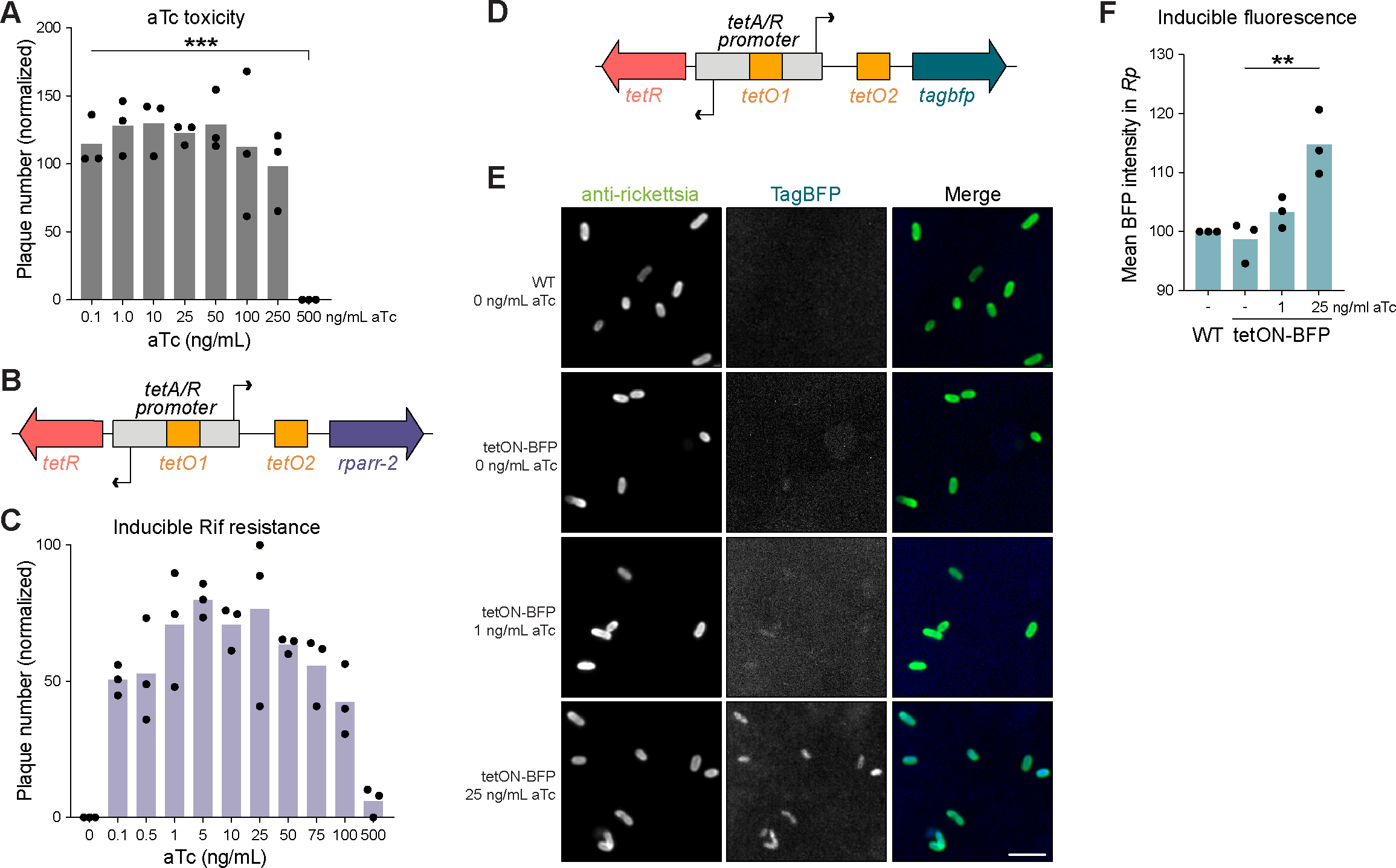
The Tet-On system enables conditional gene expression in *R. parkeri*. **(A)** Anhydrotetracycline (aTc) toxicity curve in *R. parkeri*. Plaque assays were performed on Vero cell monolayers with varying concentrations of aTc indicated. The number of plaques formed at each aTc concentration was normalized to the no aTc control for each independent experiment (n = 3). *** represents p < 0.001 by ordinary one-way ANOVA with *post hoc* Tukey’s test. **(B)** Schematic of the Tet-On system cloned into pRAM18dSGA. The tet repressor, TetR, binds two tet operator sites (*tetO*) to block gene expression in the absence of aTc. The *rparr-2* gene, which confers resistance to rifampicin, was placed under the control of Tet-On. Diagram not drawn to scale. **(C)** aTc induction of rifampicin resistance. Varying concentrations of aTc were added 30 mpi during plaque assays in Vero host cell monolayers. Each well shown had rifampicin added (200 ng/mL final concentration). All conditions shown were normalized to a no aTc and no rifampicin control well per independent experiment (n = 3). **(D)** Schematic of *tagbfp* cloned into the Tet-On system. The *tagbfp* gene was codon optimized for expression in *R. conorii*^14^. Diagram not drawn to scale. (**E & F**) aTc induction of TagBFP during infection. A549 cell monolayers were infected with *R. parkeri* harboring a plasmid containing *tagbfp* under the control of Tet-On. aTc was added 24 hpi, then samples were fixed at 48 hpi and subsequently imaged. (D) All images were set to the same minimum and maximum grey values per channel for comparison of BFP intensity. Scale bar, 2 µm. (E) Blue fluorescence from the expression of *tagbfp* was quantified for each bacterium across three independent experiments. ** denotes p < 0.01 using an ordinary one-way ANOVA.

We introduced the Tet-On system into *R. parkeri* by cloning the *tetA/R* bidirectional promoter into a pRAM18-based plasmid^13^ (Fig. 1B). This bidirectional promoter drives the expression of the tet repressor (*tetR*) gene and a downstream gene of interest (*tetA*) in the reverse and forward directions, respectively. The promoter region contains two tet operator (*tetO*) sites, to which TetR binds to block expression in both directions. Upon binding of a tetracycline derivative like aTc, TetR undergoes a conformational change that causes it to release the *tetO* binding site, thereby allowing gene expression from the *tetA/R* promoter^29^. To test inducible expression from this pRAM-based Tet-On system, we cloned the rifampicin resistance gene *rparr-2*^34^ under the control of the forward *tetA* promoter (Fig. 1B). We then challenged an *R. parkeri* strain harboring this plasmid with rifampicin and varying concentrations of aTc and monitored plaque formation over five days. In the absence of aTc, the strain is sensitive to rifampicin treatment as expected, with no plaques observed in Vero cell monolayers (Fig. 1C). In contrast, plaques were formed upon induction with aTc at concentrations as low as 0.1 ng/mL. The number of plaques trended upward with increasing concentrations of aTc, peaking between 1-25 ng/mL aTc. At 50 ng/mL aTc and above, we noticed a downward trend in the number of plaques formed, likely due to the additive toxic effects of rifampicin and high concentrations of aTc. These results show that the Tet-On system enables tunable and inducible expression of *rparr-2* in *R. parkeri* with minimal leakiness.

Because plaque formation is an endpoint assay with a population-level readout, we sought to examine how inducible expression with the Tet-On system varied across individual bacteria. Therefore, we cloned the blue fluorescent protein TagBFP (*tagbfp*) under the control of the pRAM-based Tet-On system (Fig. 1D). We then carried out a 48 h infection of A549 cells, inducing with aTc at 24 hpi. With this 24 h induction, we detected dose-dependent increases in BFP signal at 1 and 25 ng/mL aTc (Fig. 1E, F). Furthermore, we observed consistent levels of BFP signal across individual bacterial cells (Fig. 1E), indicating that the Tet-On system can be used to conditionally express genes of interest uniformly throughout the population. Still, BFP expression was modest even at the highest induction condition, only increasing in pixel intensity by ∼15% relative to WT *R. parkeri* lacking a BFP gene (Fig. 1F). In an attempt to increase BFP expression, we tested various concentrations of aTc above 25 ng/mL and reduced the induction time to 12 hours to mitigate any toxic effects from high doses of aTc. However, we were unable to significantly increase BFP expression, even at aTc levels as high as 2000 ng/mL (Supplementary Fig. 1). This result suggested that the promoter, even if fully activated, does not give rise to high levels of gene expression. As an alternative approach, we attempted to increase BFP expression upon induction by engineering *tetO* sites into strong promoters from *R. parkeri*, like P*_ompA_* and P*_ompB_*(Supplementary Fig. 2). Indeed, the intensity of BFP fluorescence with these engineered promoters was drastically higher relative to the original Tet-On system (Supplementary Fig. 2). The inducible P*_ompA_* and P*_ompB_* systems exhibited ∼100% and ∼1,000% increases, respectively, in mean pixel intensity relative to WT *R. parkeri* lacking *tagbfp*. However, at the population level, BFP expression was starkly bimodal with only ∼40% of the population strongly expressing BFP and the rest appearing BFP-negative. As a point of comparison, we generated a strain with the same *tetO*-engineered P*_ompA_* but lacking *tetR*, which therefore expresses *tagbfp* constitutively, and this strain displayed a ∼490% increase in pixel intensity relative to WT (Supplementary Fig. 2). The bimodality seen in the P*_ompA_*and P*_ompB_* inducible systems was not observed in the constitutive version of the *tetO*-engineered P*_ompA_*, meaning that the bimodal expression is not inherent to the modified promoter. While further optimization will be required to develop inducible promoters with high levels of uniform gene expression, these data demonstrate the feasibility of conditional expression using the Tet-On system in *R. parkeri*.

### Expression of dCas9 in Rickettsia parkeri via Tet-On

The lack of scalable and efficient methods to perform targeted genetic manipulation has been a major hurdle toward understanding the molecular details of the rickettsial infectious lifecycle. To address this issue and further expand the rickettsial genetic toolkit, we set out to develop a system for targeted gene knockdown in *R. parkeri*. Given its ease of use and successful application in numerous bacterial species, we chose CRISPRi^20,21^ as a candidate method for targeted genetic knockdown in *R. parkeri*. After numerous failed attempts to clone the CRISPRi components under the control of constitutive promoters on the pRAM18 backbone (data not shown), we decided to use our Tet-On inducible promoter system to express dCas9 and the constitutive promoter P*_rpsL_* to express the gRNA (Fig. 2A), similar to what was done in *Caulobacter crescentus*^25^, another member of Alphaproteobacteria. Despite the low expression of *tagbfp*, we hypothesized that this system might be ideal for CRISPRi given that lower levels of dCas9 expression are better tolerated and sufficient for gene knockdown in other bacteria^19,35,36^.

**Figure 2.**
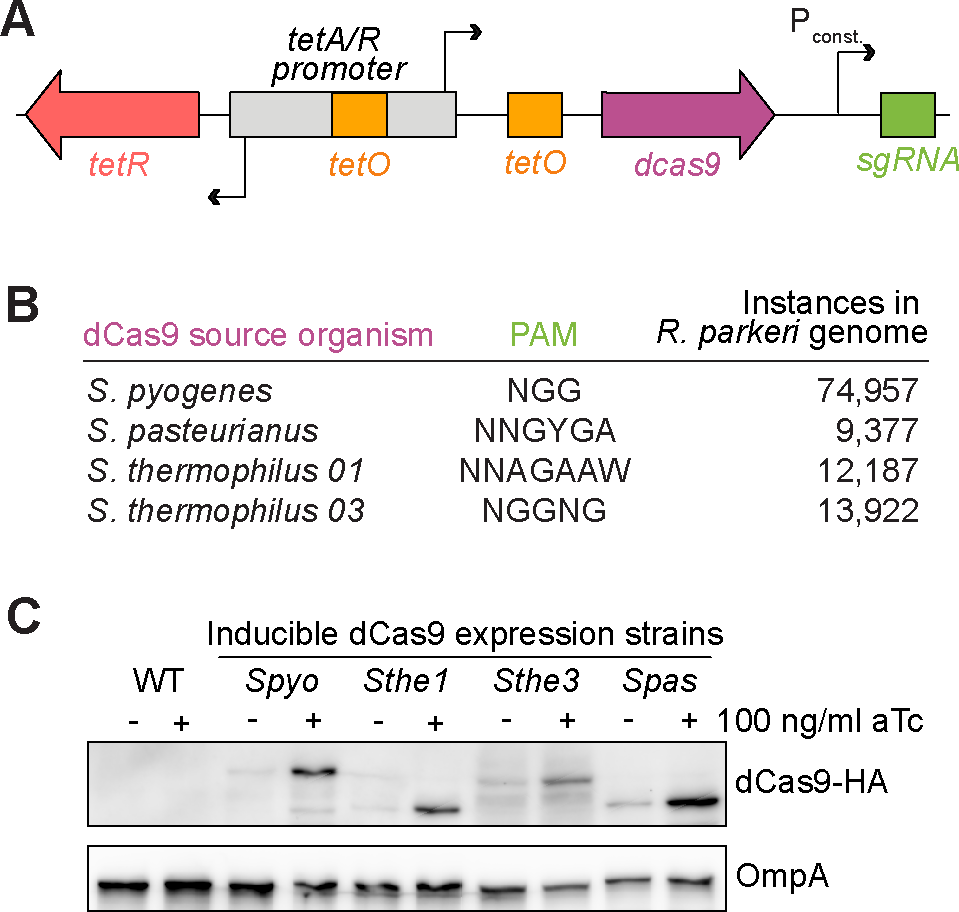
pRAM18-Tet-On can be used to express dCas9 in *R. parkeri*. **(A)** Schematic of pRAM18-based CRISPRi system. Expression of dCas9 is driven by the Tet-On promoter and the sgRNA is driven by the constitutive promoter P*_rpsL_*. **(B)** Four dCas9 variants were cloned into pRAM18dSGA. Each dCas9 variant recognizes a distinct PAM, with each PAM found in varying instances in the *R. parkeri* genome. **(C)** Expression of dCas9 in *R. parkeri*. Each dCas9 variant was tagged with a C-terminal HA epitope and expression -/+ aTc was visualized by Western blot, as well as OmpA as a loading control.

Since dCas9 proteins require binding to a protospacer adjacent motif (PAM) to license DNA binding at the target sequence^22^, the PAM limits the sequence space that is targetable via CRISPRi. Therefore, we reasoned that by selecting four candidate dCas9 variants that recognize different PAMs (Fig. 2B), we could maximize our chances of successful expression and expand the targetable range within the rickettsial genome. Two of these variants, *Streptococcus pasteurianus* (*Spas*) and *Streptococcus thermophilus* 03 (*Sthe3*), were successfully used in *C. crescentus*^25^. In addition, we tested the dCas9s derived from *Streptococcus pyogenes* (*Spyo*), which has been commonly used in many other bacteria^19^, and *S. thermophilus* 01 (*Sthe1*), which was successfully used for CRISPRi in *M. tuberculosis*^24^. Based on the different PAMs of these dCas9s (Fig. 2B), there are 96,521 targetable sites in total in the *R. parkeri* genome, making much of the 1.3 Mb genome targetable if all four dCas9s were functional.

We next needed to determine if *R. parkeri* could express each of these dCas9 variants during infection. To this end, we generated strains harboring each variant of dCas9, to which we appended a C-terminal HA tag as previously described^37^, and infected A549 cells for 72 h. We induced expression with 100 ng/mL aTc for the last 24 h of infection before harvesting cell lysates for Western blot analysis. We observed successful expression of all four dCas9 variants by Western blot (Fig. 2C). Each strain had elevated levels of dCas9 upon aTc induction, but also displayed leaky expression in the uninduced condition. Altogether, these data demonstrate successful expression of dCas9 in *R. parkeri* during infection of human cells using the Tet-On system.

### CRISPRi can be used to knockdown rifampicin resistance in R. parkeri

Given the successful expression of dCas9 during infection, we wanted to determine if CRISPRi could be used to knock down gene expression in *R. parkeri*. We chose to target *rparr-2* as it allows easy measurement of knockdown efficiency by quantifying sensitivity to rifampicin (Fig. 3A). Because we needed to test four dCas9 variants with different PAMs and the optimal parameters for selecting gRNA target sites in *R. parkeri* were unknown, we modified the original *rpsL* promoter region of the *rparr-2* locus in the *Himar1 mariner*-based transposon-containing plasmid pMW1650^10^ by enriching all four PAMs in the 100 bp immediately upstream of the predicted transcription start site. We introduced this test locus into the *R. parkeri* chromosome via random transposon insertion into the *ompA* locus, which has been documented to be dispensable during mammalian infection^17^. We performed a plaque assay on Vero host cells using this transposon integrant and determined that it forms plaques like the wild-type parental strain, as expected (data not shown).

**Figure 3.**
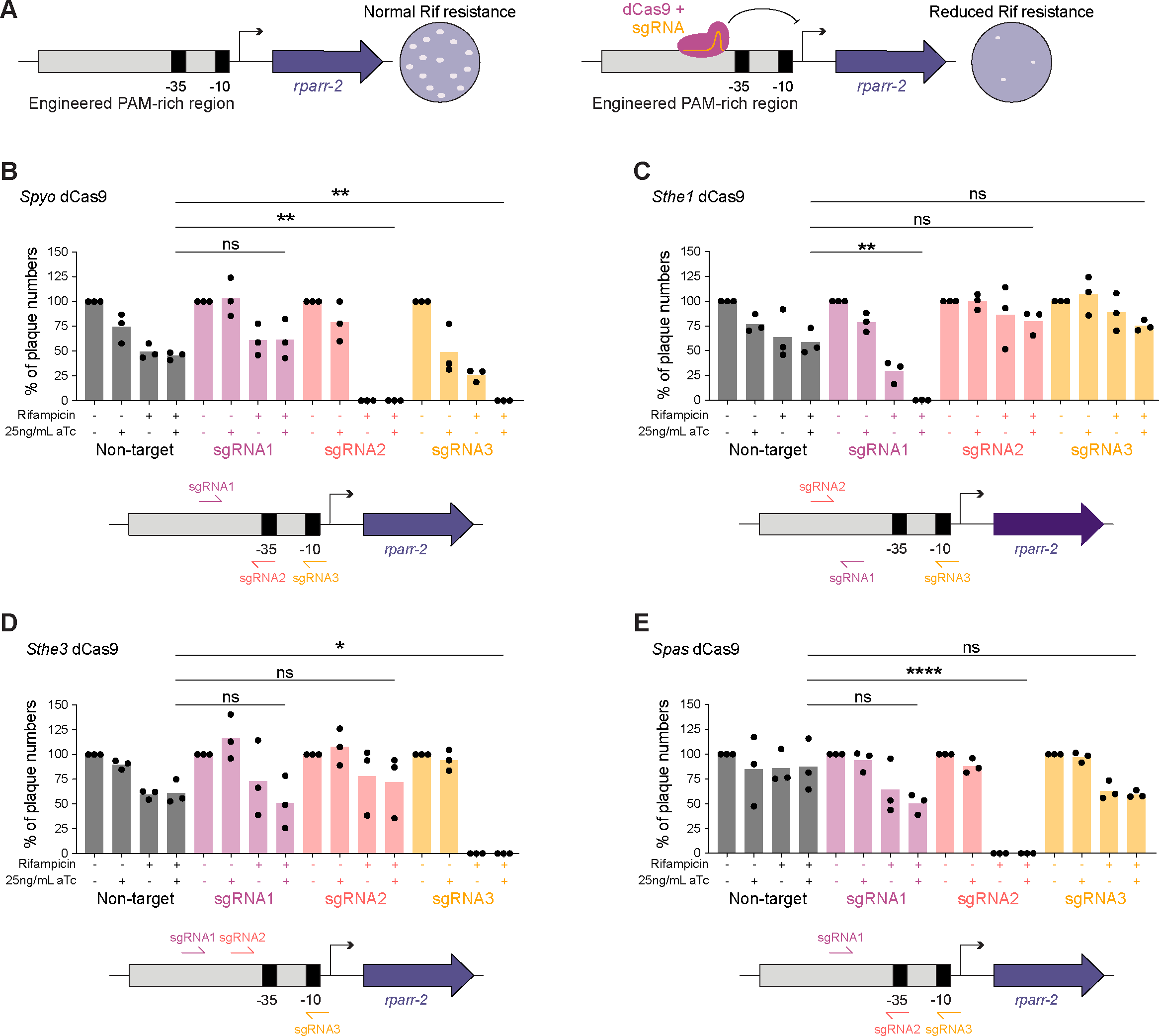
CRISPRi knockdown of a rifampicin resistance gene. **(A)** Schematic of the engineered locus and screen to test knockdown of rifampicin resistance. The *rpsL* promoter driving expression of *rparr-2* in the pMW1650 plasmid was modified to include additional PAMs to allow for testing of various dCas9 variants. Successful CRISPRi-mediated knockdown *rparr-2* would sensitize strains to treatment with rifampicin, while strains with nonfunctional CRISPRi would remain resistant to rifampicin. Spectinomycin selection ensures that the strains maintain the plasmid encoding the CRISPRi components. (**B**-**E**) Quantification of CRISPRi-mediated knockdown of rifampicin resistance via plaque assay. A549 cell monolayers were infected with *R. parkeri* strains encoding the *S. pyogenes* dCas9 (B), *S. thermophilus* 01 dCas9 (C), *S. thermophilus* 03 dCas9 (D), *S. pasteurianus* dCas9 (E). For each dCas9 variant and sgRNA combination, the same volume of *R. parkeri* stock was added to each well, and then the number of plaques was normalized to the no aTc and no rifampicin condition for a total of n = 3 independent experiments. Statistical significance was determined by ordinary one-way ANOVA with *post hoc* Tukey’s test (* denotes p < 0.05, ** denotes p < 0.005, **** denotes p < 0.0001). Schematics below each bar graph depict the relative locations of each sgRNA tested for each dCas9.

We next introduced the four dCas9s into this strain harboring the *rparr-2* insertion. For each dCas9, we designed three different gRNAs targeting the promoter region of the *rparr-2* test locus at varying distances from the promoter, covering both the template and nontemplate strands (Fig. 3). In this experimental setup, successful CRISPRi-mediated gene knockdown would sensitize the strain to rifampicin upon induction with aTc. In contrast, strains would remain resistant to rifampicin if CRISPRi knockdown failed (Fig. 3A). We tested each strain by monitoring plaque formation in Vero host cell monolayers 5 dpi. Remarkably, all four dCas9s tested successfully knocked down *rparr-2* expression, as observed by a decrease in the number of plaques formed relative to the non-target (NT) control gRNA plus aTc (Fig. 3B-E). For *Spyo* dCas9, two of the three guides tested yielded knockdown of rifampicin resistance (Fig. 3B). The remaining three dCas9s each had one successful gRNA out of the three guides tested (Fig. 3C-E). However, the majority of the gRNAs that exhibited successful knockdown of *rparr-2* showed a reduction in plaque number both in the induced and uninduced conditions, indicating leakiness of the system. Two gRNAs, *Spyo* gRNA3 and *Sthe1* gRNA1 showed only a partial reduction in plaques in the uninduced condition, compared to zero plaques observed in the induced condition, suggesting partial inducibility with these dCas9/gRNA combinations. Interestingly, all of the guides that yielded significant knockdown of *rparr-2* targeted the nontemplate strand, similar to what was observed in other systems including *C. crescentus*^25^. While further optimization will be required to decrease the leakiness of the inducible promoter system, our results demonstrate CRISPRi-mediated targeted gene knockdown for the first time in *Rickettsia*.

### CRISPRi knockdown of the rickettsial virulence factor Sca2

Because the knockdown experiments described above targeted an exogenously introduced locus with an engineered promoter, we wanted to test the ability of our CRISPRi system to target endogenously encoded virulence factors in the *R. parkeri* genome. We chose to target the *sca2* gene, which encodes a formin-like actin nucleator that mediates long actin tail formation (Fig. 4A)^11,38,39^. *Sca2* was an ideal target for several reasons: (1) the transposon mutant of *sca2* has been well characterized^11,39^; (2) loss of actin tail formation is easily observable via fluorescence microscopy; and (3) *sca2* does not appear to be encoded in an operon, making targeting with CRISPRi simpler.

**Figure 4.**
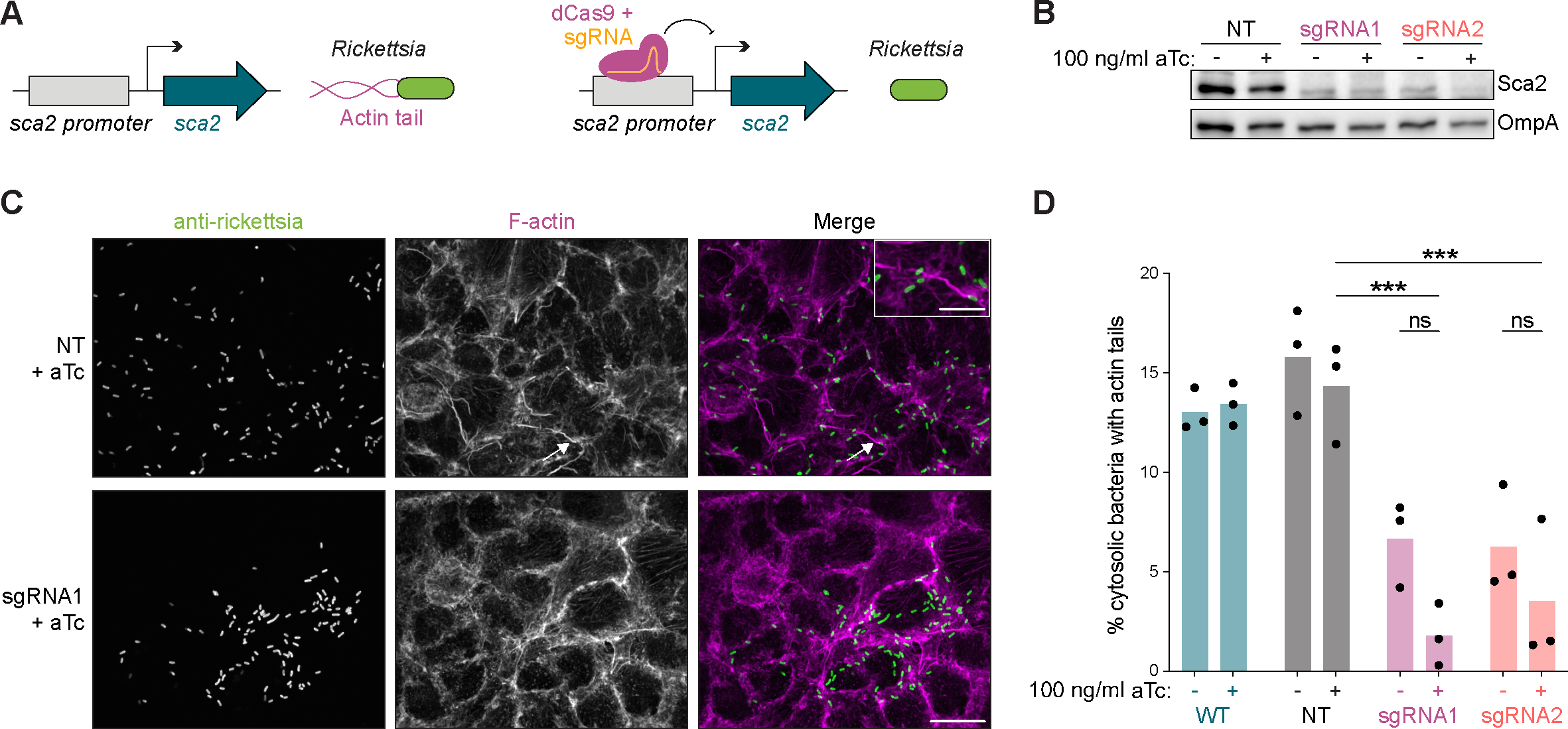
CRISPRi knockdown of the rickettsial virulence factor *sca2*. **(A)** Schematic of *sca2* knockdown experiment. Sca2 is a formin-like actin nucleator responsible for forming long actin tails during *R. parkeri* infection. CRISPRi-mediated knockdown of *sca2* should result in decreased actin tail formation. **(B)** CRISPRi targeting leads to decreased expression of Sca2 protein. A549 host cell monolayers were infected with *R. parkeri*. aTc was added to infections 48 hpi and lysates were harvested at 72 hpi. Sca2 and OmpA (loading control) protein levels were visualized via Western blotting. (**C** & **D**) Measurement of actin tail formation by immunofluorescence. A549 cell monolayers were infected with *R. parkeri* for 28 h, with aTc being added to appropriate wells at the time of infection. These samples were then fixed, stained, and imaged to visualize (C) and quantify (D) actin tail formation. The white arrow indicates an actin tail, which is shown in greater detail in the inset. Scale bar, 10 µm and 5 µm in inset. For each condition, at least 300 bacteria were quantified in each of n = 3 independent experiments. *** denotes p < 0.001, determined by one-way ANOVA with *post hoc* Tukey’s test.

Based on our results from the knockdown of rifampicin resistance, we designed two gRNAs (gRNAs 1 and 2) for *Spas* dCas9, targeting the nontemplate strand upstream of the coding region in the endogenous *sca2* locus. We first tested if we could detect obvious loss of *sca2* expression using CRISPRi. We harvested *R. parkeri* infections of A549 host cells at 3 dpi, with 100 ng/mL aTc induction in the last 24 h before sample collection. Indeed, we were able to detect a decrease in Sca2 protein levels in both gRNA1 and gRNA2 relative to the NT control (Fig. 4B).

We then tested if CRISPRi-mediated silencing of *sca2* also led to the expected reduction in actin tail frequency^11,39,40^. We infected A549 host cells with strains harboring each of the gRNAs, with and without aTc, and fixed samples at 28 hpi for subsequent immunofluorescent staining and confocal microscopy (Fig. 4C). As predicted from our Western blot data, both gRNA1 and gRNA2 resulted in a significant decrease in tail formation (Fig. 4D), corroborating successful knockdown of *sca2*. Similar to the results from the experiments targeting *rparr-2*, no significant difference was observed between induced and uninduced, indicating leakiness of the inducible promoter system expressing dCas9. Taken together with the results above, our experiments demonstrate successful knockdown of the endogenously encoded *sca2* virulence factor in *R. parkeri* using CRISPRi.

## DISCUSSION

The *Rickettsia* genus is comprised of obligate intracellular bacteria and several members are neglected and emerging human pathogens^41^. Understanding their unique biology and mechanisms of pathogenesis has been hindered by the paucity of genetic tools available to carry out functional-genetic studies. Here, we have expanded the genetic toolkit by developing systems for conditional gene expression and targeted gene knockdown in rickettsiae. These tools will be invaluable for dissecting the genetic and molecular requirements of the rickettsial infectious life cycle and revealing novel biology at the host-pathogen interface.

The ability to control gene expression with a small molecule inducer will be a powerful tool for performing mechanistic investigations of key rickettsial virulence genes. For instance, by varying the timing and dosage of induction, this tool will allow us to study the kinetic requirements of a given virulence gene during the infectious life cycle. While conditional expression systems have been developed in *C. burnetii* and *C. trachomatis*, many obligate intracellular bacteria still lack them^42^. Here, we have adapted the Tet-On system for use in *R. parkeri*, enabling conditional gene expression for the first time in a *Rickettsia* species. While this system uses aTc as the inducer molecule, our work suggests that other small molecule inducers could also potentially be implemented in rickettsiae, like IPTG or arabinose. Further development of orthogonal conditional expression systems will open even more possibilities for genetic studies in rickettsiae.

Inducible promoter systems also offer other valuable applications including controlled expression of toxic proteins and conditional expression of other genetic tools like CRISPRi. In fact, heterologous expression of Cas9 and derivatives like dCas9 have been found to have toxic effects in various other bacteria, including model organisms like *E. coli*^19,36^. We were unable to clone dCas9 under a constitutive promoter into the pRAM18dSGA backbone in *E. coli*, which we speculate was due to dCas9 toxicity. Using the Tet-On system to control the expression of dCas9, we were able to successfully demonstrate CRISPRi-mediated targeted gene knockdown for the first time in the Rickettsiales order.

CRISPRi makes performing targeted gene knockdowns relatively simple as one only needs to design guide RNA sequences that target the locus of interest. However, the target space of CRISPRi is limited to sites within the genome with the appropriate PAM adjacent to the gRNA target sequence. Each dCas9 variant has a unique PAM that is recognized. Therefore, it is useful to have multiple variants of dCas9 to choose from to expand the range of targetable sites in the genome. Similar to previous reports^43^, we also observed some flexibility in the *S. pasteurianus* dCas9 PAM requirements, with successful knockdown arising from gRNAs using both the NNGTGA and NNGCGA PAMs (e.g. sgRNA1 and sgRNA2, respectively, from the *sca2* KD assay). In our study, we tested four variants of dCas9 that recognize different PAMs. Given previous work that found varying success with different dCas9 variants in different bacteria^24,25^, we were surprised that all four of the dCas9 variants we tested yielded successful gene knockdown. This variety of functional dCas9s affords us an expanded range of targetable sites within the rickettsial genome.

The design of optimal gRNA sequences also depends on other factors, including strand biases and proximity to the promoter. Similar to what was observed in several other bacteria, including the alphaproteobacterium *C. crescentus*^25^, we found a stark preference for gRNAs targeting the nontemplate strand. Of the 14 gRNAs tested in this study, 9 of the gRNAs were complementary to the nontemplate strand. Incredibly, 7 of these 9 gRNAs targeting the nontemplate strand yielded significant gene knockdown. Most of the gRNAs were designed to target near the promoter, ideally within 100 bp of the predicted transcription start site of the target gene. Expanded testing of gRNAs targeting other rickettsial genes will be required to determine the precise parameters for optimal gRNA design in rickettsiae.

For both the inducible promoter and CRISPRi systems, there are limitations in their practical use in their current state. For example, expression of *bfp* was difficult to detect in the presence of aTc, suggesting weak expression of certain genes in the “on” state. We attempted to overcome this by engineering *tetO* sites into known strong rickettsial promoters like P*_ompA_* and P*_ompB_*, but this yielded a starkly bimodal distribution of BFP expression (high BFP fluorescence vs. no BFP signal) across the bacterial population. Given that these experiments were performed in the presence of antibiotic selection to maintain the plasmid, we do not believe this is due to plasmid loss. However, we cannot completely rule out the possibility of plasmid instability as a cause, and the observed distribution could be due to plasmid rearrangements or variability in plasmid copy number. Interestingly, similarly uneven expression from a Tet-On system was also observed in *C. trachomatis*, which was attributed to variable metabolic states at certain time points of infection during the chlamydial life cycle^32^. We also attempted to implement the same tet-inducible promoter used in *C. trachomatis*^32^, but this promoter was not functional in *R. parkeri* (data not shown). Additional work will be required to better understand what underlies the bimodal distribution we observe with our Tet-On system in *R. parkeri*.

Fortunately, despite the weak expression of *bfp,* the Tet-On system allows for sufficient expression of dCas9 for gene knockdown. It is possible that the low expression from our Tet-On system might be necessary to avoid toxicity in rickettsiae, as dCas9 has been shown to be toxic when expressed at high levels in other bacteria^19^. Despite the Tet-On system displaying no detectable leakiness when controlling the expression of *rparr-2* and *bfp*, the expression of *dcas9* is less tightly controlled in the absence of aTc, with some variants showing more leaky expression than others. Notably, we observed some amount of inducibility for two of the gRNAs that yielded knockdown of *rparr-2* (*Spyo* gRNA3 and *Sthe1* gRNA1). This variability in leakiness between different gRNAs is similar to what was observed in *M. smegmatis*, where they had to build and optimize a collection of new inducible promoters to enable targeting of essential genes^24^. We tested the optimized promoter that was implemented in *M. tuberculosis*^24^, but it demonstrated weak and leaky expression in *R. parkeri* (data not shown). Alternatively, it might be possible to decrease the leakiness of the system by also placing the gRNA under the control of a separate inducible promoter^19^. Beyond optimization of the inducible promoter system itself, it might also be possible to improve the inducibility of CRISPRi by introducing mismatches into non-seed regions of gRNAs to weaken their affinities to their respective target sites, similar to what has been done in other systems^44,45^. Another alternative approach could be to express a Cas9-specific anti-CRISPR protein to antagonize dCas9 in the absence of inducer, but at a low enough level so that the anti-CRISPR could be overcome during induction of dCas9 expression^46^.

Nevertheless, the tools presented here open new roads for detailed investigations into the biology and pathogenesis of these important human pathogens. CRISPRi provides a platform for efficient and scalable targeted gene knockdown in rickettsiae, opening the possibility to directly probe the *in vivo* relevance of rickettsial genes that had previously only been studied through biochemical or exogenous expression assays. Our CRISPRi platform, combined with future improvements in transformation efficiency in rickettsiae, could allow for the first large-scale reverse genetic screens in *Rickettsia*. Moreover, sequence-specific binding of dCas9 can be used for other technological applications, such as CRISPR activation (CRISPRa) to increase expression of endogenous loci^20^. Overall, our work introduces two new methods for controlling gene expression in rickettsiae, which will ultimately be critical for gaining new insights into fundamental host-pathogen interactions and understanding how these neglected and emerging pathogens cause disease.

## MATERIALS AND METHODS

### Cell culture

Vero African green monkey kidney epithelial and A549 human lung epithelial cell lines were obtained from the University of California, Berkeley Cell Culture Facility (Berkeley, California). Vero cells were maintained in Dulbecco’s modified Eagle’s medium (DMEM; Gibco catalog number 11965118) containing 5% fetal bovine serum (FBS). A549 cells were maintained in DMEM with 10% FBS. Assays measuring *tagbfp* expression, *dcas9* expression, *rparr-2* knockdown, and *sca2* knockdown were conducted using DMEM containing tetracycline-negative FBS. Cell lines were confirmed to be mycoplasma-negative in a MycoAlert PLUS assay (Lonza catalog number LT07-710) performed by the Koch Institute High-Throughput Sciences Facility (Cambridge, Massachusetts).

### Plasmid construction

pRL0027 was generated from pRAM18dSGA[MCS]^13^ (kindly provided by Ulrike Munderloh) by removing *gfp* and changing the promoter of the spectinomycin resistance cassette to P*_rpsL_*with an *ompA* terminator proceeding the gene. pRL0081 was generated from pRL0027 by cloning the Tet-On promoter from pdCas9-bacteria (Addgene catalog number 44249) along with *rparr-2* and *gfpuv* under the control of Tet-ON with a single *ompA* terminator sequence. pRL0117, pRL0234, pRL0235, pRL0236, and pRL0237 were all generated similarly to pRL0081 but cloned downstream of the Tet-On promoter was a codon-optimized version of *tagbfp* for *R. conorii*^14^, *S. pyogenes dCas9* with constitutively expressed *gRNA-SapI*, *S. thermophilus 01 dCas9* with constitutively expressed *gRNA-SapI*, *S. thermophilus 03 dCas9* with constitutively expressed *gRNA-SapI*, and *S. pasteurianus dCas9* with constitutively expressed *gRNA-SapI*, respectively. A C-terminal HA tag was appended to all dCas9 variants. pRL0200 was cloned via gene synthesis (Twist Biosciences) using the *tagbfp* sequence from pRL0117 and the *ompA* promoter from *R. parkeri* with a *tetO1* operator sequence from pRL0081. pRL0203 was generated similarly to pRL0200 but with additional synthetic DNA fragments including the *tetR* gene with its promoter and *gfpuv* from pRL0081. pRL0202 was generated similarly to pRL0203 but with codon-optimized *tagbfp* under the control of a synthetic promoter constructed by adding two *tetO2* sequences from pRL0081 to the *ompB* promoter from *R. parkeri*.

pRL0057 was generated from pMW1650^10^ by replacing the *rpsL* promoter region with a 100 bp sequence that was modified to include additional PAM sequences.

Guide RNA plasmids were cloned via restriction cloning by digesting the pRL0234, pRL0235, pRL0236, and pRL0237 backbones with SapI and gel purifying the cut vector. Short oligonucleotides (Sigma) were designed to have compatible overhangs upon annealing and were ligated into the cut pRL0234-pRL0237 backbones. gRNA sequences were manually selected based on proximity and position relative to the predicted transcription start site and likelihood of off-target effects. Potential off-target sites for each gRNA were screened for using Cas-OFFinder^47^ (http://www.rgenome.net/cas-offinder/) and a modified Python script based on a previously published package^48^. A full list of gRNA sequences is provided in Supplementary Table 1.

### Generation of *R. parkeri* strains

Wild type *R. parkeri* strain Portsmouth (kindly provided by Chris Paddock) and all derivatives were propagated by infection and mechanical disruption of Vero cells grown in DMEM containing 2% FBS at 33°C as previously described^14^. These bacterial stocks were further purified using 2 µm syringe filtering (Whatman) as previously described^40^. Bacteria were clonally isolated from plaques formed from Vero host cell monolayer infection in the presence of agarose overlays as previously described^8^. All bacterial stocks were stored as aliquots at -80°C in brain heart infusion media (BHI; Fisher Scientific, DF0037-17-8) to minimize freeze-thaw cycles. Titers were measured via plaque assay on Vero cells and quantified at 5 dpi.

Plasmids were introduced into *R. parkeri* via small-scale electroporation as previously described^8^ with approximately 1 µg of dialyzed plasmid DNA. Selection was started 24 h after electroporation by overlaying a mixture of 0.5% agarose, DMEM with 2% FBS, and either rifampicin (200 ng/mL final concentration) or spectinomycin (50 µg/mL). The sites of transposon insertions for generating the modified *rparr-2*-containing strain for testing dCas9 knockdown were determined by semi-random nested PCR and Sanger sequencing as previously described^8^.

### Plaque assays

Plaque assays were conducted as previously described^8^. Briefly, confluent Vero cell monolayers grown in 6-well plates were washed in PBS and subsequently infected with *R. parkeri* in a humidified chamber and rocked for 30 min at 37°C. DMEM with 2% FBS and 0.5% agarose was overlaid on top of the infected cells, and this was incubated in a humidified chamber at 33°C with 5% CO_2_ for 5 d. Plaque assays were then imaged and analyzed using Fiji/ImageJ. For assays involving aTc induction, a small volume of concentrated aTc solution was added on top of the molten agarose overlays to give the appropriate final aTc concentration.

### BFP expression assays

Confluent A549 host cell monolayers were grown on 12-mm coverslips in 24-well plates and infected at a multiplicity of infection (MOI) of approximately 0.05. *R. parkeri* was added to the media and centrifuged at 200 x g for 5 min at room temperature (RT). These infections were subsequently incubated at 33°C and anhydrotetracycline (aTc) was added to appropriate wells at 24 hpi. Samples were then fixed at 48 hpi by adding 4% paraformaldehyde in phosphate-buffered saline (PBS) for 10 min at RT. Fixed samples were then washed in PBS and residual paraformaldehyde was quenched by incubating samples with 0.1 M glycine for 10 min at RT. Next, samples were washed with PBS and incubated with blocking buffer (2% bovine serum albumin [BSA] in PBS) for 30 min at RT. Samples were treated with primary and secondary antibodies suspended in blocking buffer for 1 h each, with three PBS washes after each incubation. Phalloidin conjugated to Alexa Fluor 647 (Invitrogen catalog # A22287) was used to detect actin and mouse anti-*Rickettsia* 14-13 (kindly provided by Ted Hackstadt) was used to detect *R. parkeri*. Coverslips were mounted with ProLong Gold Antifade mountant (Invitrogen catalog # P36934). For each condition, at least 300 bacteria were imaged using a 100× UPlanSApo (1.35 NA) objective. Images were processed with Fiji/ImageJ. CellProfiler^49^ was used to measure blue fluorescence intensity within the bounds of individual bacteria as detected by anti-*Rickettsia* staining.

### Actin tail assay

Confluent A549 host cell monolayers were infected and processed similarly to above with minor modifications. Infections were carried out at an MOI of approximately 0.1 to 0.5. Before infection, the media was replaced with fresh DMEM including appropriate antibiotics and anhydrotetracycline (100 ng/mL final concentration) in appropriate wells. Infected samples were fixed with paraformaldehyde at 28 hpi. Hoechst stain (Invitrogen catalog # H3570) was used to detect host cell nuclei. Image analysis was performed with Fiji/ImageJ. For every replicate of each strain and condition, at least 3 fields of view and at least 300 bacteria were analyzed to calculate the percentage of cytosolic bacteria with actin tails (>1 bacterial length). This was performed in triplicate.

### Immunoblotting of Sca2 and dCas9-HA from infected host cell lysates

Fresh DMEM including appropriate antibiotics was added to confluent A549 cell monolayers, which were subsequently infected with strains of *R. parkeri* harboring plasmids with *sca2* or *dcas9-ha* under the control of the Tet-On promoter. aTc was added 48 hpi. Then at 72 hpi, the infected A549 host cell monolayers were resuspended in loading buffer (50 mM Tris-HCl [pH 6.8], 2% sodium dodecyl sulfate [SDS], 10% glycerol, 0.1% bromophenol blue, 5% β-mercaptoethanol) and boiled for 20 minutes with vortexing. These samples were analyzed via Western blotting using rabbit anti-Sca2 (kindly provided by Matthew Welch), mouse anti-HA (BioLegend catalog number 901501), and mouse anti-OmpA 13-3 (kindly provided by Ted Hackstadt).

### Statistical analyses

All statistical analyses were performed using GraphPad Prism 10. Graphical representations, statistical parameters, and significance are noted in figure legends. Statistical significance was defined as *p* < 0.05.

**Supplementary Figure 1. Expression of TagBFP from Tet-On using higher concentrations of aTc.**

A549 cell monolayers were infected with *R. parkeri* harboring a plasmid containing Tet-On::*tagbfp*. aTc was added 16 hpi and samples were fixed at 28 hpi. 12 h induction was used to minimize toxic effects from high concentrations of aTc. Fixed samples were imaged using spinning disk confocal fluorescent microscopy. All images were set to the same minimum and maximum grey values per channel for comparison of BFP intensity. Scale bar, 5 µm.

**Supplementary Figure 2. Expression of TagBFP from engineered aTc-responsive rickettsial promoters.**

(Left) A549 cell monolayers were infected with *R. parkeri* harboring a plasmid that expressed *tagbfp* from various promoters. The strong rickettsial promoters P*_ompA_* and P*_ompB_* were engineered to be aTc-responsive by adding *tetO* sites into the promoters. aTc was added 4 hpi and samples were fixed at 28 hpi. The samples were subsequently imaged via spinning disk confocal fluorescent microscopy. All images were set to the same minimum and maximum grey values per channel for comparison of BFP intensity. Red arrow indicates bacterium with no detectable *tagbfp* expression, blue arrowhead indicates bacterium expressing *tagbfp*. Scale bar, 10 µm. (Right) Schematic of rickettsial promoters engineered to be aTc-inducible. Diagrams not drawn to scale.

## ACKNOWLEDGMENTS

We thank Ulrike Munderloh, Matthew Welch, Michael Laub, Chris Paddock, and Ted Hackstadt for sharing strains and reagents. We are grateful to members of the Lamason laboratory for helpful discussions. Work in the Lamason laboratory is supported in part by the National Institutes of Health (R01 AI155489) and by the Office of the Assistant Secretary of Defense for Health Affairs through the Tick-Borne Disease Research Program (TB200032). Opinions, interpretations, conclusions, and recommendations are those of the authors and are not necessarily endorsed by the Department of Defense. JM is a Damon Runyon Fellow supported by the Damon Runyon Cancer Research Foundation (DRG-2396-20). DLE was supported by the MIT Undergraduate Research Opportunities Program (UROP) Office, including through the Peter J Eloranta Summer Undergraduate Research Fellowship, the John Reed UROP Fund, and the Joseph Woo (1988) UROP Fund for the Life Sciences.

